# FLIM-based pH measurements reveal incretin-induced rejuvenation of aged insulin secretory granules

**DOI:** 10.1101/174391

**Authors:** Martin Neukam, Anke Sönmez, Michele Solimena

## Abstract

Insulin is stored in dense-core secretory granules (SGs) and is released from beta cells in two distinct phases upon glucose stimulation. Newly synthesized insulin SGs are secreted preferentially, but the underlying mechanism of this phenomenon remains elusive. The relationship of SG age with their intraluminal pH is of particular interest: proinsulin conversion by prohormone convertases follows the acidification of immature SGs by the vacuolar proton-translocating ATPase (v-ATPase). v-ATPases may also participate in the formation of the fusion pore for SG exocytosis, with intraluminal alkalinization inhibiting membrane fusion. Previous studies examined the luminal pH of SGs on a population level only. Here we measured the pH-dependent lifetime changes of eCFP fused to the ICA512-RESP18 homology domain to assess for the first time the luminal pH of individual age-defined SGs in insulinoma INS-1 cells by fluorescence-lifetime imaging microscopy. We show that 2-4-hour-old young SGs have a pH of ~5.5, while 26-28-hour-old SGs have a pH of ~6.2. Remarkably, the GLP-1 receptor agonist Exendin-4 prompted the re-acidification of old SGs in a glutamate-dependent fashion, while it did not affect the pH of young SGs. This study demonstrates that insulin SGs change their pH over time - a change that is reversible by insulin secretagogues. Hence, it provides novel insight into the mechanisms accounting for aging and exocytosis of SGs and suggests that their ‘rejuvenation’ may be exploited to enhance insulin secretion in diabetes.

## Introduction

The hormone insulin is synthesized and secreted by pancreatic beta cells. It is the key regulator of glucose uptake and therefore it is essential for glucose homeostasis in vertebrates. Insulin is stored within dense-core secretory granules (SGs) and is released from beta cells in two distinct phases upon elevation of circulating glucose levels (*Curry et al., 1968*). The first phase lasts only few minutes and is characterized by a burst of insulin secretion, while the second, prolonged phase lasts until glycemia returns to fasting levels. Hallmarks of type 2 diabetes, which is the most common form of the disease, are impaired glucose-stimulated insulin secretion, especially during the first phase; increased proinsulin release at disease onset; and perturbation of the pulsatile pattern of insulin release in the basal state (*Porte, 1991*). Understanding the general principles that govern insulin SG exocytosis and turnover is therefore of paramount importance to elucidate the pathogenesis of diabetes (*Müller et al, 2017b*).

According to the prevalent view, insulin is stored in two distinct pools of SGs. During the first phase, insulin secretion results from the exocytosis of readily releasable SGs that are already docked to the plasma membrane prior to glucose-induced depolarization of the beta cell plasma membrane (*Rorsman and Renström, 2003*). The second phase is instead mainly mediated by the recruitment to the plasma membrane of more distant SGs from the so-called reserve pool. This model has been revised with the identification of a set of SGs, termed „restless newcomers‟ that upon entering the field of TIRF microscopy are recruited for fusion with the plasma membrane without any time delay (*Shibasaki et al., 2007*). As restless newcomers contributed to both phases of insulin secretion, while newly synthesized SGs are preferentially secreted (*Schatz et al., 1975; Gold et al., 1982, Halban et al., 1982*) it is of great interest to correlate the age of SGs to their motility and properties in general.

To this aim, our laboratory recently developed a protocol for the unequivocal labeling and imaging of age-distinct SGs using a Ins-SNAP reporter (*Ivanova et al., 2013*). Exploitation of this approach enabled us to formally validate the preferential release of newly synthesized insulin SGs ex-vivo and reveal that young SGs display greater motility compared to their older counterparts (*Ivanova et al., 2013; Hoboth et al., 2015*). Specifically, we found that ageing of SGs entails their reduced competence for microtubule-mediated transport, while F-actin marks aged SGs destined for intracellular degradation within multigranular bodies/lysosomes via autophagy (*Müller et al., 2017a; Müller et al, 2017b*). However, the reasons why SGs decrease their motility and secretory competence over time and are then targeted to autophagy remain unknown.

Membrane traffic of vesicles requires a strict control of their luminal pH. For example, KDEL-receptor mediated retrieval of ER-resident proteins from the Golgi back to the ER is a process that takes advantage of the different intraluminal pH between these two compartments along the secretory pathway (*Wilson et al., 1993*). In general, it is well established that the vesicular luminal pH progressively drops between the ER and the plasma membrane (*Wu et al., 2001*). Proteolytic processing of prohormones initiates upon acidification of post-Golgi immature SGs, which leads to the activation of prohormone convertases (*Schmidt et al., 1995*). For proinsulin, a pH of 5.5 is required for its optimal conversion to insulin (*Orci et al., 1986*). The acidification of SGs depends on the activity of the vacuolar proton-translocating ATPase (v-ATPase), which is also responsible for the acidification of other compartments, such as lysosomes, endosomes, synaptic vesicles and the Golgi complex (*Kane 2006, Forgac et al., 2007*). Increasing evidence suggests that v-ATPases are not only critical for the maturation of SGs, but also for their ability to undergo exocytosis, possibly by participating in the formation of the fusion pore (*Hiesinger et al., 2005; Strasser et al., 2011*). Notably, intraluminal alkalinization may induce conformational changes in the cytosolic domain of the v-ATPase, thereby inhibiting SG exocytosis (*Poëa-Guyon et al., 2013b*), and/or the organization of the surrounding microfilaments (*Vitavska et al, 2003; Vitavska et al, 2005*). In view of these considerations, we aimed therefore to re-assess the pH on individual SGs. Our findings show that the luminal pH of SGs correlates with their age and that the alkalinization of old SGs can be reverted in a glutamate-dependent fashion by treatment with Exendin-4, which is an insulin secretagogue related to the incretin hormone GLP-1.

## Material and Methods

### Plasmids and materials

RESP18-HD-CFP was cloned as previously described (*Torrko et al., 2015*). Antibodies against insulin (mouse monoclonal, Sigma #I-2018), EEA1 (mouse monoclonal, Thermo #MA5-14794) and LAMP2 (rabbit polyclonal, Thermo #PA1-655) were used at dilution 1:200. Similarly, secondary anti-mouse/rabbit Alexa^568^ antibodies were used. Calibration buffers consisted of 15 mM MES, 15 mM HEPES, 140 mM KCl and 10 µM Nigericin (Tocris, 5 min) and calibrated to pH 5.0 to 7.5 in 0.5 intervals. Bafilomycin A1 (Sigma, 60 min) was used at 100 nM, DIDS (Tocris, 60 min) at 10 µM and IAA-94 (Enzo LifeScience, 60 min) at 10 µM, Exendin-4 (Sigma, 15 min) at 50 nM, Rose Bengal (Sigma, 60 min) at 30 µM and DM-glutamate (Sigma, 15 min) at 1 mM. All drugs were supplemented to standard culture medium with 11 mM glucose prior to imaging.

### Reagents and Cell culture

Rat insulinoma INS-1 cells, which were a kind gift of Dr. Claes Wollheim (Geneve, Switzerland), were grown on high precision coverslips for localization studies and in 35 mM MatTek glass bottom dishes for live cell imaging. The cells were cultured in standard INS-1 cell medium with 11 mM glucose and used 3-4 days post-transfection. For co-localization studies, cells were fixed with 4% PFA (20 min, RT), permeabilized with Triton X-100 (for immunostaining of insulin) or 0.1% saponin (for immunostainings of EEA1 and LAMP2), blocked and stained with antibodies. Live cell fluorescence-lifetime imaging microscopy (FLIM) was performed in a temperature (37°C) and CO_2_ (5%) controlled chamber. Insulin-SNAP was labelled as previously described (*Ivanova et al., 2013, Hoboth et al., 2015*). Briefly, the medium of INS-1 cells was changed to include either non-fluorescent SNAP-Cell Block (1:400) or SNAP-Cell TMR-Star (1:750). After 30 min the medium was aspirated, the cells washed twice with PBS and replaced by non-modified medium. The cells were washed three times for 30 min.

### Microscopy and data analysis

Co-localization analysis were performed by super-resolution microscopy with a DeltaVision OMX SIM (GE Healthcare), using an Olympus Plan ApochromatN 60x oil objective with an NA of 1.42. Stacks with a Z step size of 125 nm were acquired and images reconstructed using the SoftWoRx software (SoftWoRx, Germiston, South Africa). Confocal and FLIM microscopy was done on a Zeiss LSM780 confocal microscope with a Becker&Hickl dual-channel TCSPC (SPC-152). A Zeiss alpha Plan-Apochromat 63x oil objective with an NA of 1.46 was used. First, a confocal image was taken using Zen2012 and then switched to the FLIM mode. Cells were excited with a 440 nm laser at 50 Hz and imaged until ~2000 photons were collected in the brightest pixel. For calibration the medium was changed to one of the calibration buffers 5 min prior to imaging. Image analysis was performed using FIJI and SPCImage. Mean values and confidence interval (CI) were calculated using Excel.

## Results & Discussion

### Generation of a genetic reporter for insulin SG pH

Previous measurement of the luminal pH were performed using (non-)ratiometric dyes that accumulate in acidic compartments. Using this approach, it was suggested that the SG pH becomes more alkaline upon glucose stimulation (*Barg et al., 2001, Eto et al., 2003*). Stiernet et al (2006), in contrary, reported acidification upon glucose treatment. Despite being the most commonly used technique, ratiometric dyes lack specificity for the SG population and can therefore accumulate also in other organelles, such as lysosomes or the Golgi complex. A more targeted approach was the utilization of an eGFP-based construct, which showed acidification of the SG lumen after glucose stimulation and alkalinization upon cell treatment with IBMX (*Tompkins et al., 2002*). However, it is difficult to normalize for expression levels and therefore only relative pH changes were recorded. Hence, a reliable SG pH reporter should be specifically targeted to SGs and accurately report the luminal pH in the range from 5 to 7.5.

To meet these requirements, we fused the enhanced cyan fluorescent protein (CFP) to the RESP18 homology domain (RESP18-HD) of ICA512. CFP has been shown to be a reliable marker for pH measurements by FLIM (*Poëa-Guyon et al., 2013a*), whereas the RESP18-HD was shown to contain information that are necessary and sufficient for the sorting of ICA512 and other cargoes to the SGs (*Torkko et al, 2015*). In transfected INS-1 cells fixed with PFA, RESP18-HD-CFP was enriched in punctate structures also positive for endogenous insulin (Fig. 1A), but negative for the early endosomal or lysosomal markers EEA1 and LAMP2, respectively (Fig. S1). The fluorescence lifetime signal of RESP18-HD-CFP was calibrated by incubating nigericin-permeabilized INS-1 cells in MES-HEPES buffers which vary in their pH from 5.0 to 7.5 (Fig. 1B). The calibration curve indicated that the fluorescent lifetime of the reporter was linear within this pH range (Fig. 1C). Interestingly, the lifetime of RESP18-HD-CFP^+^ SGs appeared to be heterogeneous (Fig. S2).

**Figure 1.**
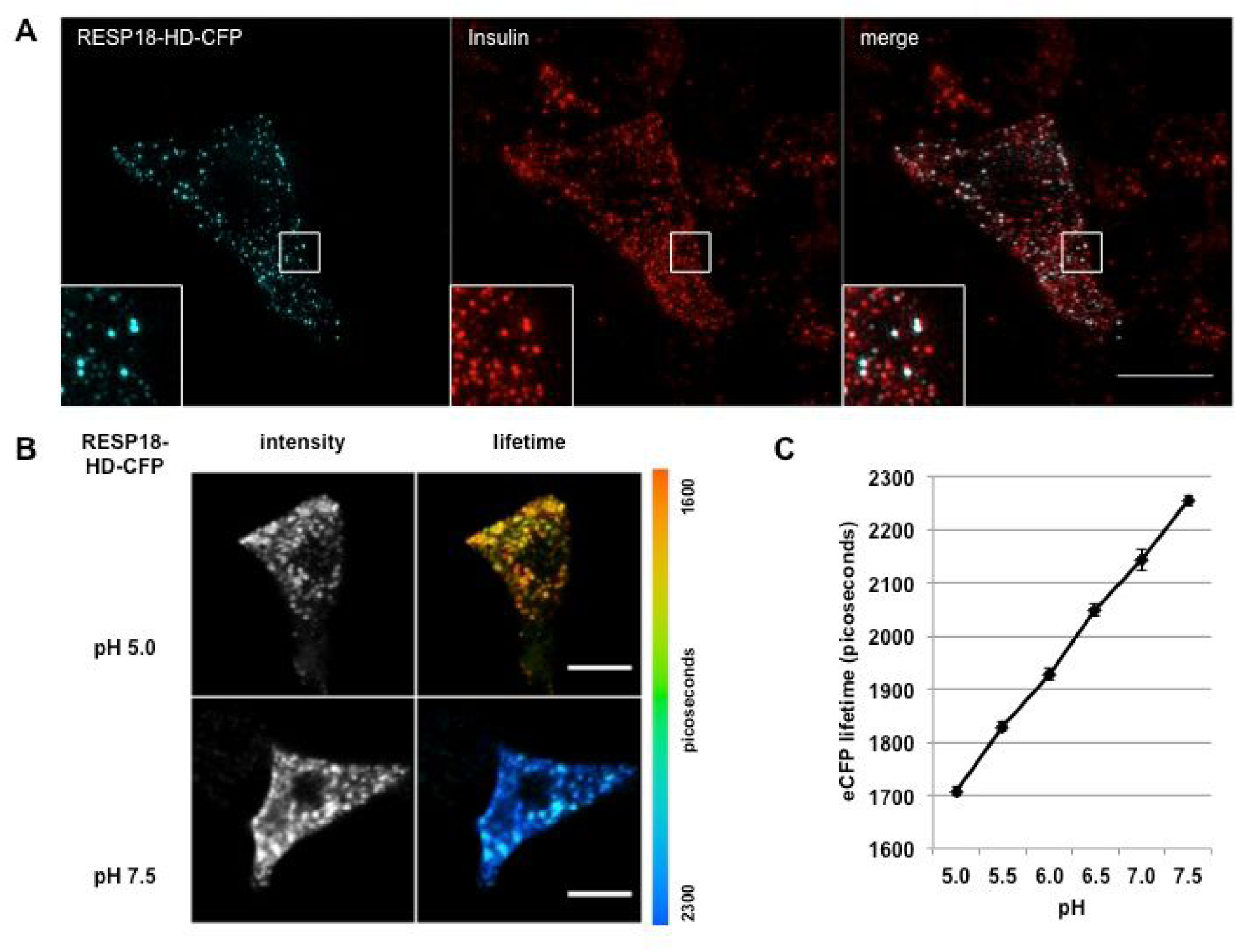
Characterization of the SG pH reporter. (**a**) Imaging by super-resolution microscopy for transiently transfected RESP18-HD-CFP and endogenous insulin in INS-1 cells. The inset is a 8.53 magnification of the cell area within the smaller square. Scale bar: 10 µm. (**b**) Representative images acquired by confocal microscopy for calibration of the fluorescence-lifetimes of CFP in INS-1 cells which were transiently transfected with RESP18-HD-CFP and permeabilized with 10 µM nigericin in buffers having the desired pH between 5.0 and 7.5. Images were acquired 5 min after treatment. Acidic pH correlates with a short lifetime of CFP and is depicted in orange. At neutral pH the lifetime is longer and is depicted in blue. Scale bar: 10 µm. (**c**) Calibration curve of the CFP fluorescence lifetimes corresponding to (**b**).

### SG change their pH over time

Since an acidic pH was postulated to affect the release probability of chromaffin SGs (*Poëa-Guyon et al., 2013b*) and we found that in INS-1 cells 4-6-hour-old insulin SGs are preferentially released in comparison to 24-26-hour-old insulin SGs (*Ivanova et al., 2013*), we investigated a potential correlation between pH and SG age (Fig. 2A). Co-transfection of RESP18-HD-CFP with the Ins-SNAP reporter for SG age enabled us to assess that the luminal pH of 2-4-hour-old SGs (n=110 in 26 cells) was 5.49±0.09 (Fig. 2B), which corresponds to the optimal pH for the activity of the prohormone convertases that process proinsulin (*Hutton, 1982*). Up to 10-hour-old (young) RESP-18-HD-CFP^+^ SGs were acidic with a pH of 5.49±0.09. Remarkably, however, their pH became more alkalinize with time, being 6.21±0.10 in 26-28-hour-old-SGs (n=127 in 38 cells). Yet, treatment with the v-ATPase inhibitor Bafilomycin A1 changed the luminal pH of both 2-4-hour-old SGs (n=49 in 11 cells) and 26-28-hour-old-SGs (n=96 in 16 cells) to 6.99±0.14 and 6.96±0.06, respectively. These findings validate that age-distinct SG pools depend on the v-ATPase activity for their luminal acidification, but that over time the pH of SGs shifts to more alkaline values.

**Figure 2.**
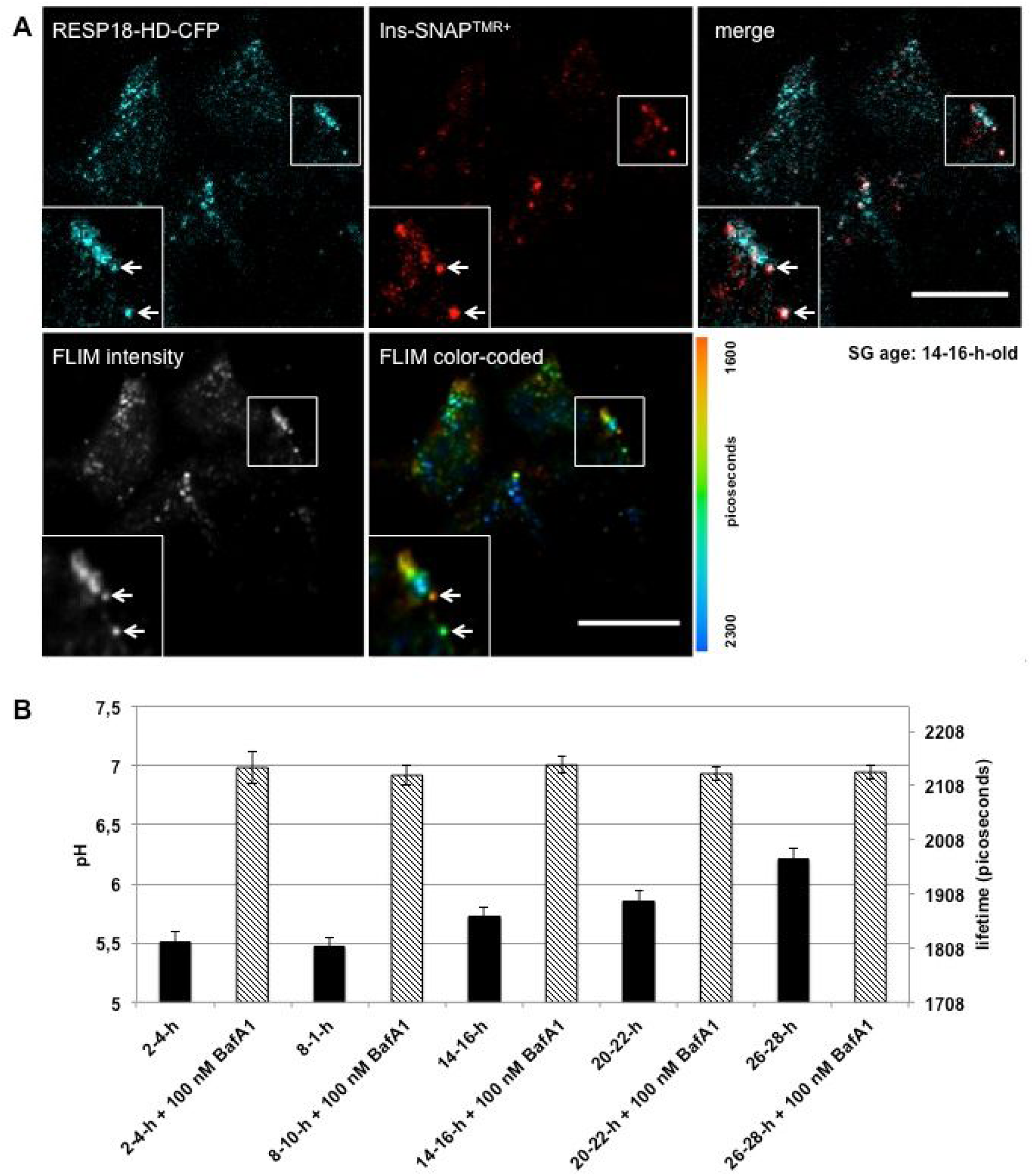
Correlation of SG pH and age. (**a**) Representative confocal images for determination of SG pH and age in living INS-1 cells transiently co-transfected with the RESP-18-HD-CFP and Ins-SNAP reporters. Upper panel: Confocal microscopy imaging of RESP-18-HD-CFP^+^ SGs having a defined age (in this case 14-16-hour-old) based on the fluorescence signal of the age reporter Ins-SNAP labeled with TMR. The inset is a 5.12 magnification of the cell area within the smaller square. Lower panel: FLIM for the measurement of the pH in the same age-defined SGs based on the lifetime of CFP. Inset (same inset area as in (**a**)): two adjacent 14-16-hour-old RESP-18-HD-CFP^+^, Ins-SNAP^TMR+^ SGs (arrows) display different lifetimes, i.e. their luminal pH being either 5.5 (orange) or 6.2 (green). The nearby RESP-18-HD-CFP^+^ puncta likely correspond to SGs younger or older than 14-16 hour-old SGs, and therefore negative for Ins-SNAP^TMR+^. Scale bar: 10 µm. (**b**) Quantification of the pH in age-distinct SGs pools based on images similar to (**a**) in cells untreated or treated with 100 nM Bafilomycin A1 (BafA1) for 30 min, which inhibits the v-ATPase. Images were acquired immediately after treatment. Data are represented as mean ± CI of 95%.

### Exendin-4 but not chloride affects SG pH

Fine-tuning of the luminal pH is generally considered to be achieved via either of two pathways. Counterion conductance can be a limiting factor for proton translocation across membranes as unfavourable electro-chemical gradients are reached. Alternatively, leakage pathways could release protons back to the cytosol and thereby fine-tune the pH. Chloride is known as a counterion in the re-acidification of endosomes and previous reports suggested the presence of a chloride channel on insulin SGs (*Barg et al., 2001, Deriy et al., 2009*), however this remains debated (*Maritzen et al., 2008, Jentsch et al., 2010*). We found that exposure of INS-1 cells to the chloride channel blockers indanyloxyacetic acid (IAA-94) or 4,4'-diisothiocyanato-stilbene-2,2'-disulphonic acid (DIDS) did not affect the pH of 2–4-hour-old SGs and 26–28-hour-old SGs (Fig. 3A and B).

**Figure 3.**
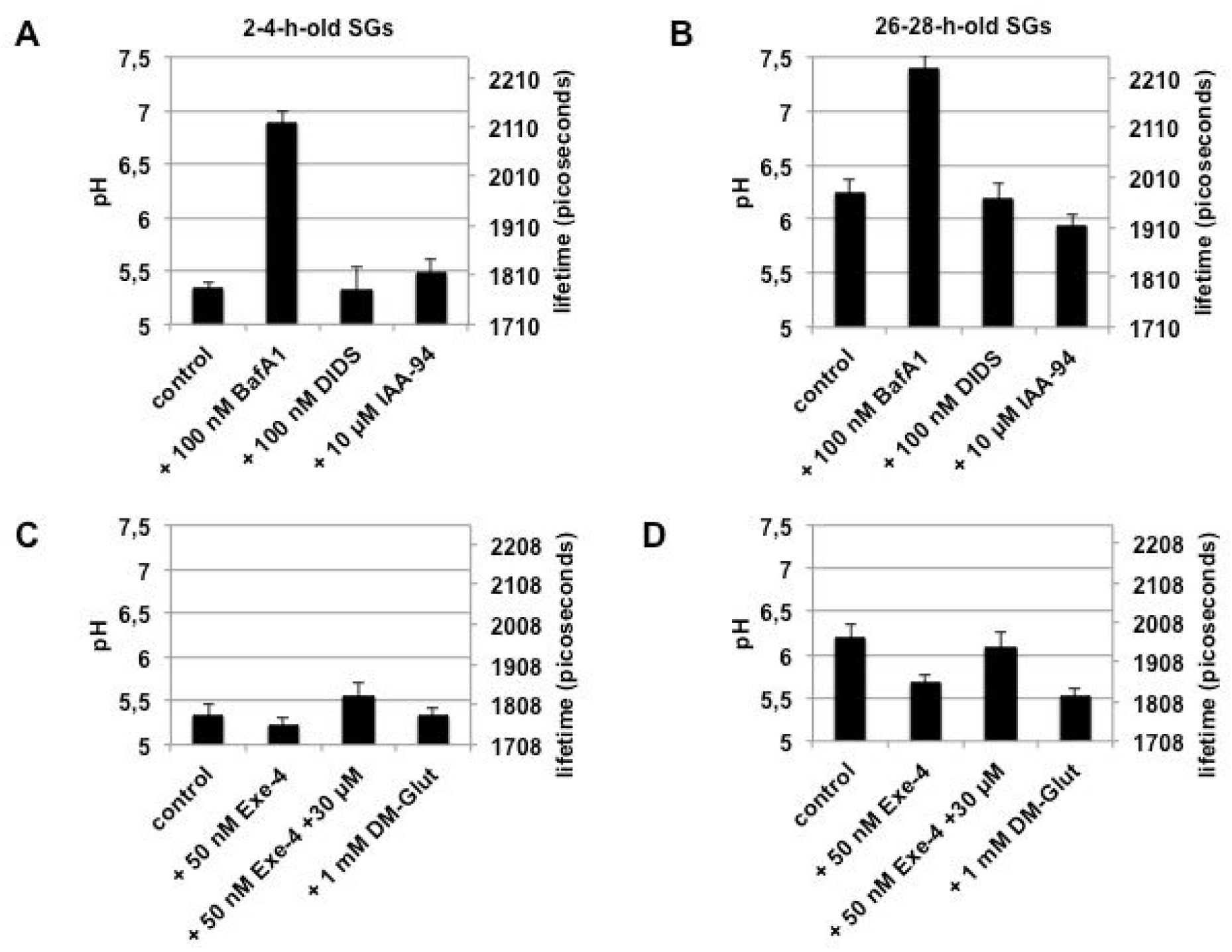
Impact of treatments on the pH of age-distinct SGs. (**a-b**) Measurement of pH in 2-4-hour-old (**a**) and 26-28-hour-old (**b**) SGs in living RESP-18-HD-CFP+, Ins-SNAP^TMR+^ INS-1 cells untreated (control) or treated for 60 minutes with chloride channel inhibitors DIDS (100 nM) and IAA-94 (10 µM). (**c-d**) Measurement of pH in 2-4-hour-old (**c**) and 26-28-hour-old (**d**) SGs in living RESP-18-HD-CFP+, Ins-SNAP^TMR+^ INS-1 cells untreated (control) or treated for 15 minutes with GLP-1 receptor agonist Exendin-4 (Exe-4, 50 nM), 50 nM Exendin-4 + vGLUT inhibitor Rose Bengal (RB, 30 µM), or membrane permeable dimethyl-glutamate (DM-Glut, 1 mM). Data are represented as mean ± CI of 95%.

Glutamate, which may be involved in the amplifying pathway of glucose stimulated-insulin secretion (*Maechler, 2017*), can be uptaken into insulin SGs upon GLP-1 treatment, thereby potentiating insulin secretion (*Gheni et al., 2014*). However, it is unknown if all SGs take up glutamate and if this results in the acidification of their lumen, as previously proposed (*Maechler and Wollheim, 1999*). When INS-1 cells were treated with the GLP-1 receptor agonist Exendin-4 the luminal pH of 2-4-hour-old SGs in treated cells was unaffected relative to control cells (Fig. 3C and Tab. S1). Conversely, the luminal pH of 26-28-hour-old SGs dropped significantly, restoring the pH measured in younger SGs (Fig. 3D and Tab. S1). As suggested, glutamate uptake into SGs may be driven by the vesicular glutamate transporter vGLUT, since treatment with its inhibitor Rose Bengal prevented the Exendin-4-induced re-acidification of older SGs, while having no effect of the pH of the younger. Re-acidification of the older SGs was also elicited by treatment with the membrane-permeable dimethyl-glutamate, which however did not alter the pH of the younger SGs. We further excluded that re-acidification of the older SGs upon Exendin-4 treatment reflected their targeting to lysosomes for degradation, since 26-28-hour-old SGs were negative for the lysosomal marker LAMP2 (Fig. 4). Hence, Exendin-4-induced ‘rejuvenation’ of old SGs is likely mediated by glutamate uptake and account, at least in part, for the ability of GLP-1 to potentiate glucose-stimulated insulin secretion, and thus ameliorate glucose tolerance.

**Figure 4.**
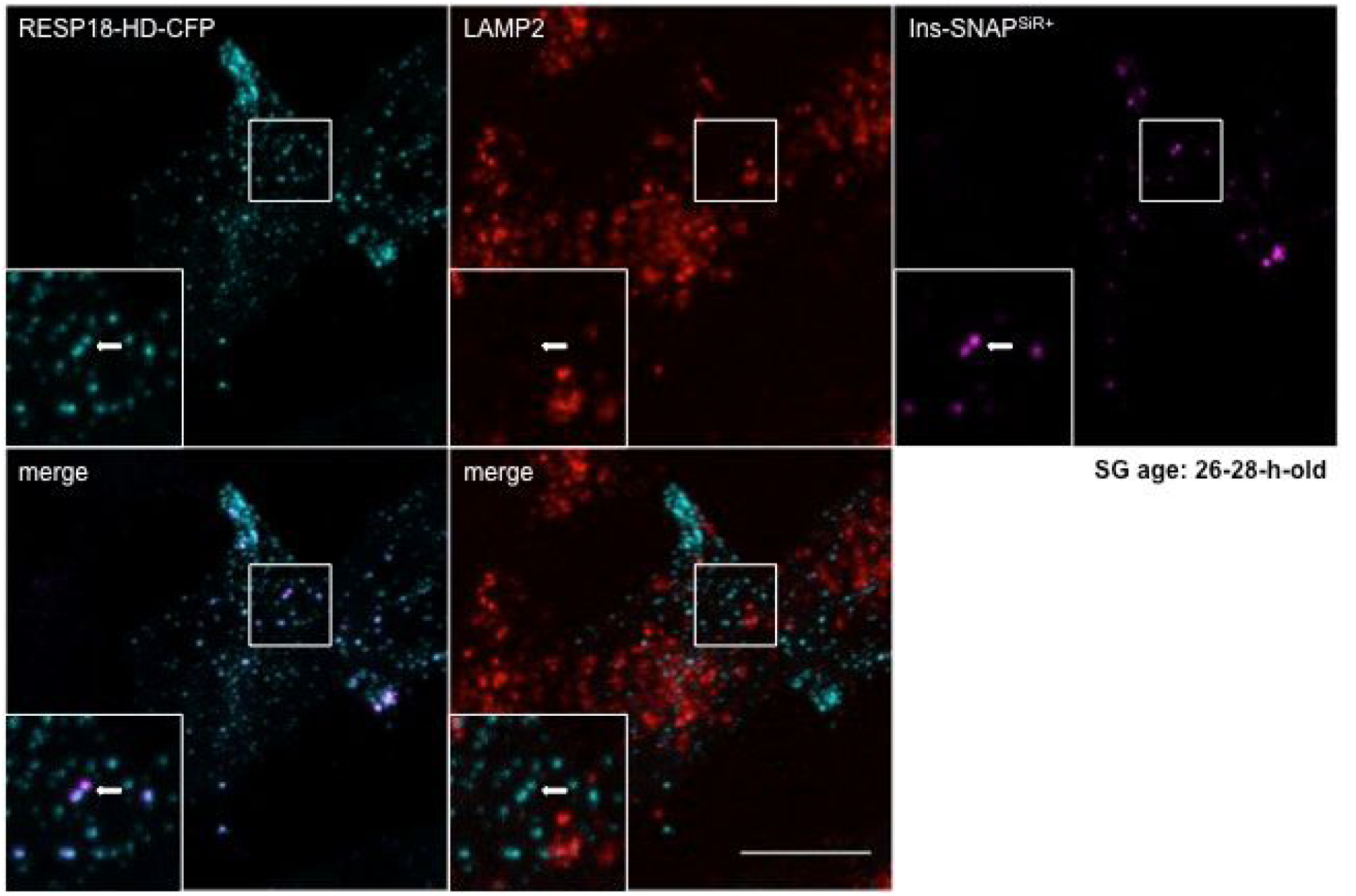
Confocal microscopy images of INS-1 cells co-transfected with the RESP18-HD-CFP and Ins-SNAP reporters and immunostained for the lysosomal marker LAMP2. Images were acquired 28 hours after labeling of cells with the fluorescent SNAP substrate SiR and 15 minutes after treatment with 50 nM Exendin-4. The inset is a 5.39 magnification of the cell area within the smaller square. The arrow points at a 26-28-hour-old Ins-SNAP^SiR+^, RESP18-HD-CFP^+^ SG, which is negative for LAMP2. Scale bar: 10 µm.

## Conclusions

Here we show that aging of SGs results in the alkalinization of their lumen, which may also hinder their ability to undergo fusion with the plasma membrane. Previously we found that aging of SGs, which reduces their competence for exocytosis, correlates with reduced microtubule-mediated transport and increased probability for disposal by autophagy. The molecular principles responsible for the change in the luminal pH of SGs over time, as for the change in SG motility, remain to be elucidated. Interestingly, however, both these changes are reversible upon treatment with Exendin-4, a major compound for the treatment of type 2 diabetes. cAMP-mediated activation of Epac2 downstream of GLP-1 signaling is associated with remodeling of the cortical actin cytoskeleton and increased mobility of SGs (*Shibasaki et al., 2007*). Our data demonstrates that GLP-1 signaling also selectively restores the luminal pH of aged SGs to 5.5, similarly to the pH present in newly-generated SGs, which are preferentially released. Whether SG pH and mobility are intimately connected remains to be ascertained, but it is tempting to speculate that a unifying process regulates both these variables, such that aged SGs in the reserve pool can be recruited back to the readily releasable pool for insulin secretion.

## Acknowledgments

We thank Bert Nitzsche for advice regarding SIM imaging, Sophie Pautot and Nicolas Vitale for advice regarding pH measurements with CFP by FLIM. We are grateful to Katja Pfriem for excellent administrative assistance. This work was supported with funds to MS by the German Center for Diabetes Research (DZD e.V.), which is financed by the German Ministry for Education and Research. MN is the recipient of a fellowship from the DFG-supported Dresden International Graduate School for Biomedicine and Bioengineering (DIGS-BB) and of a traveling fellowship from the Graduate Academy Dresden.

## Author contributions

M.N. designed and performed all the experiments. A.S. provided technical help with cell culture. M.S. conceived the project and supervised the work. The manuscript was written by M.N. and M.S.

## Footnotes

**Competing interests statement** The authors declare no competing financial interests.

## Figure and Table legends

**Figure S1.**
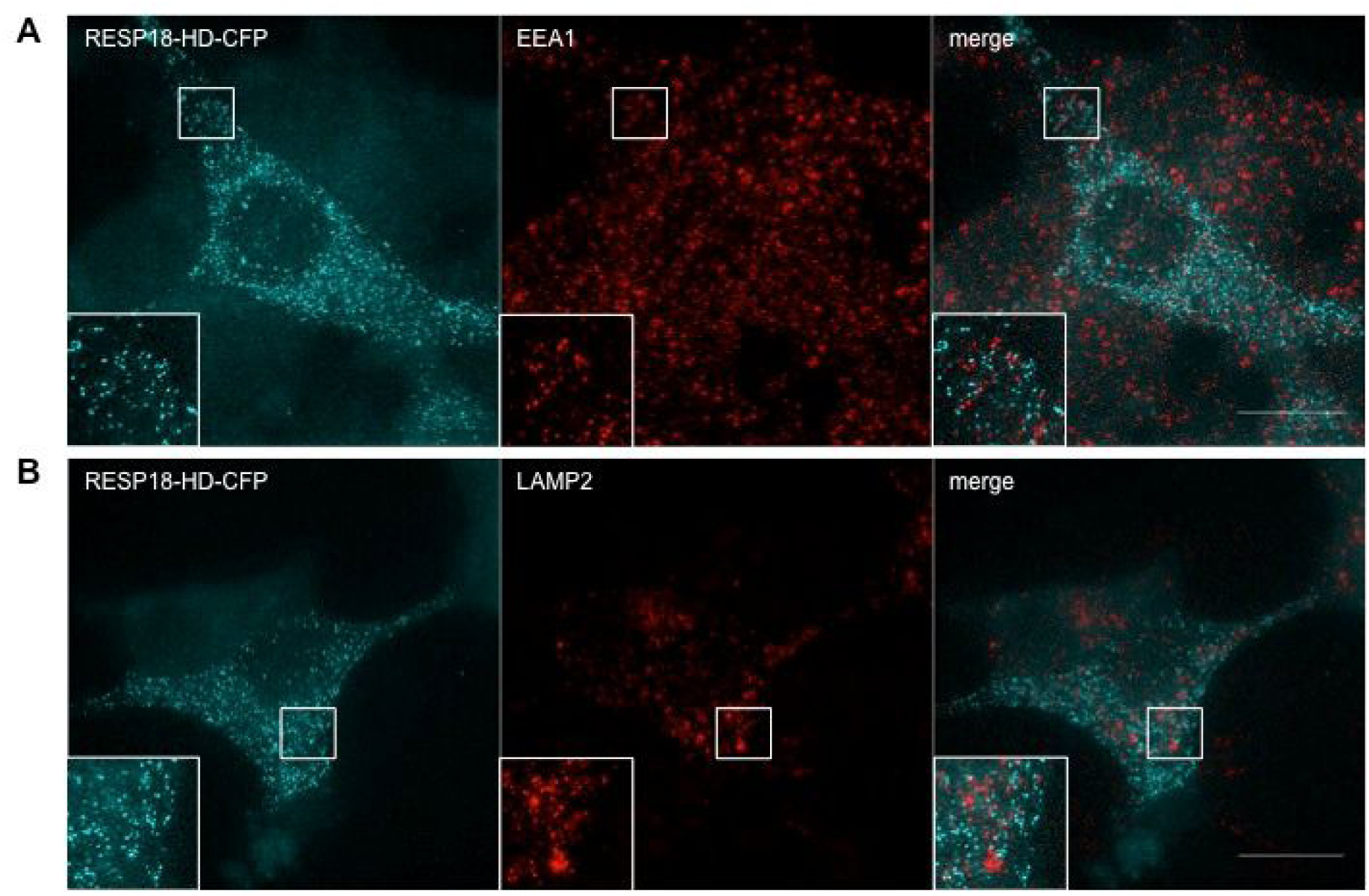
Characterization of the SG pH reporter. (**a**) Imaging by super-resolution microscopy for transiently transfected RESP18-HD-CFP and endogenous EEA1 (a) or LAMP2 (**b**) in INS-1 cells. The insets are 6.83 magnifications of the corresponding cell areas within the smaller squares. Scale bar: 10 µm.

**Figure S2.**
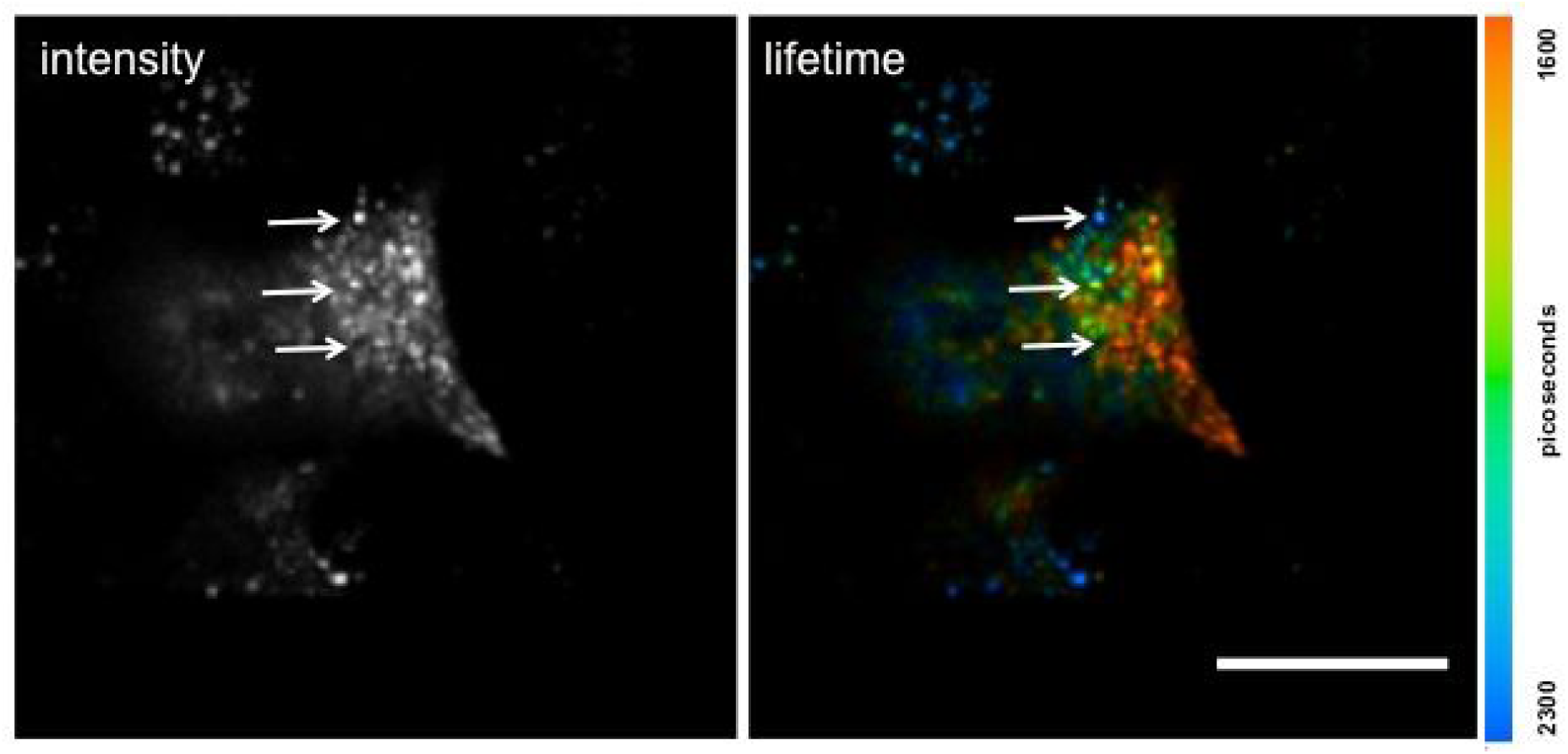
Heterogeneity of CFP fluorescence lifetime in individual SGs (arrows) of INS-1 cells transiently transfected with RESP18-HD-CFP. Scale bar: 10 µm.

**Table S1.** Effect of different treatments on SG pH. Living INS-1 cells were either kept in standard culture medium with 11.1 mM glucose or treated with Exendin-4 (50 nM), Exendin-4 (50 nM) + Rose Bengal (30 µM) or with DM-Glut (1 mM) for XX time prior to imaging by confocal-fluorescence lifetime microscopy. Data are represented as mean ± CI of 95%

|  | control | Exendin-4 | Rose Bengal | DM-Glutamate |
| --- | --- | --- | --- | --- |
| 2-4-h-old SGs | pH: $5.31 \pm 0.13$<br>(49 SGs, 11 cells) | pH: $5.20 \pm 0.10$<br>(51 SGs, 11 cells) | pH: $5.54 \pm 0.16$<br>(54 SGs, 13 cells) | pH: $5.41 \pm 0.10$<br>(91 SGs, 15 cells) |
| 26-28-h-old SGs | pH: $6.19 \pm 0.15$<br>(41 SGs, 14 cells) | pH: $5.66 \pm 0.11$<br>(71 SGs, 17 cells) | pH: $6.08 \pm 0.18$<br>(70 SGs, 18 cells) | pH: $5.51 \pm 0.10$<br>(86 SGs, 18 cells) |

## References

1. Barg, S., Huang, P., Eliasson, L., Nelson, D.J., Obermüller, S., Rorsman, P., Thévenod, F. and Renström, E. (2001). Priming of insulin granules for exocytosis by granular Cl(-) uptake and acidification. J Cell Sci, 2001, 114(Pt 11), 2145–54.

2. Curry, D.L., Bennett, L.L. and Grodsky, G.M. (1968). Dynamics of insulin secretion by the perfused rat pancreas. Endocrinology 1968, 83(3), 572–84.

3. Deriy, L.V., Gomez, E.A., Jacobson, D.A., Wang, X., Hopson, J.A., Liu, X.Y., Zhang, G., Bindokas, V.P., Philipson, L.H. and Nelson, D.J. (2009). The granular chloride channel ClC-3 is permissive for insulin secretion. Cell Metab, 2009, 10(4), 316–23.

4. Eto, K., Yamashita, T., Hirose, K., Tsubamoto, Y., Ainscow, E.K., Rutter, G.A., Kimura, S., Noda, M., Iino, M. and Kadowaki, T. (2003). Glucose metabolism and glutamate analog acutely alkalinize pH of insulin secretory vesicles of pancreatic beta-cells. Am J Physiol Endocrinol Metab 2003, 285(2): E262–71.

5. Forgac, M. (2007). Vacuolar ATPases: rotary proton pumps in physiology and pathophysiology. Nat Rev Mol Cell Biol 2007, 8(11), 917–29.

6. Gheni, G., Ogura, M., Iwasaki, M., Yokoi, N., Minami, K., Nakayama, Y., Harada, K., Hastoy, B., Wu, X., Takahashi, H., Kimura, K., Matsubara, T., Hoshikawa, R., Hatano, N., Sugawara, K., Shibasaki, T., Inagaki, N., Bamba, T., Mizoguchi, A., Fukusaki, E., Rorsman, P. and Seino, S. (2014). Glutamate acts as a key signal linking glucose metabolism to incretin/cAMP action to amplify insulin secretion. Cell Rep, 2014, 9(2), 661–73.

7. Gold, G., Gishizky, M.L. and Grodsky, G.M (1982). Evidence that glucose “marks” beta cells resulting in preferential release of newly synthesized insulin. Science 1982, 218, 56–8.

8. Halban, P.A. (1982). Differential rates of release of newly synthesized and stored insulin from pancreatic islets. Endocrinology 1982, 110(4): 1183–8.

9. Hiesinger, P.R., Fayyazuddin, A., Mehta, S.Q., Rosenmund, T., Schulze, K.L., Zhai, R.G., Verstreken, P., Cao, Y., Zhou,Y., Kunz, J. and Bellen, H.J. (2005). The v-ATPase V0 subunit a1 is required for a late step in synaptic vesicle exocytosis in Drosophila. Cell 2005, 121, 607–20.

10. Hoboth, P., Müller, A., Ivanova, A., Mziaut, H., Dehghany, J., Sönmez, A., Lachnit, M., Meyer-Hermann, M., Kalaidzidis, Y. and Solimena, M. (2015). Aged insulin granules display reduced microtubule-dependent mobility and are disposed within actin-positive multigranular bodies. Proc Natl Acad Sci U S A 2015, 112(7), E667–676.

11. Hutton, J.C. (1982). The internal pH and membrane potential of the insulin-secretory granule. Biochem J. 1982, 204(1): 171–8.

12. Ivanova, A., Kalaidzidis, Y., Dirkx, R., Sarov, M., Gerlach, M., Schroth-Diez, B., Müller, A., Liu, Y., Andree, C., Mulligan, B., Münster, C., Kurth, T., Bickle, M., Speier, S., Anastassiadis, K. and Solimena, M. (2013). Aged-dependent labeling and imaging of insulin secretory granules. Diabetes 2013, 62(11), 3687–96.

13. Jentsch, T.J., Maritzen, T., Keating, D.J., Zdebik, A.A. and Thévenod, F. (2010). ClC-3—a granular anion transporter involved in insulin secretion? Cell Metab, 2010, 12(4), 307–8.

14. Kane, P.M. (2006). The where, when, and how of organelle acidification by the yeast vacuolar H+-ATPase. Microbiol Mol Biol Rev 2006, 70(1), 177–91.

15. Maechler, P. (2017). Glutamate pathways of the beta-cell and the control of insulin secretion. Diabetes Res Clin Pract 2017, 131:149–153.

16. Maechler, P. and Wollheim, C.B. (1999). Mitochondrial glutamate acts as a messenger in glucose-induced insulin exocytosis. Nature 1999, 402(6762), 685–9.

17. Maritzen, T., Keating, D.J., Neagoe, I., Zdebik, A.A. and Jentsch, T.J. (2008). Role of the vesicular chloride transporter ClC-3 in neuroendocrine tissue. J Neurosci, 2008, 28(42), 10587–98.

a) Müller, A., Neukam, M., Ivanova, A., Sönmez, A., Münster, C., Kretschmar, S., Kalaidzidis, Y., Kurth, T., Verbavatz, J.-M. and Solimena, M. (2017). A Global Approach for Quantitative Super Resolution and Electron Microscopy on Cryo and Epoxy Sections Using Self-labeling Protein Tags. Sci Rep 2017, 7(1):23.

b) Müller, A., Mziaut, H., Neukam, M., Knoch, K.P. and Solimena, M. (2017). A 4D view on insulin secretory granule turnover in the β-cell. Diabetes Obes Metab 2017, 19 Suppl, 107–114.

20. Orci, L., Ravazzola, M., Amherdt, M., Madsen, O., Perrelet, A., Vassalli, J.D. and Anderson, R.G. (1986). Conversion of proinsulin to insulin occurs coordinately with acidification of maturing secretory vesicles. J Cell Biol 1986, 103, 2273–81.

a) Poëa-Guyon, S., Pasquier, H., Mérola, F., Morel, N. and Erard, M. (2013). The enhanced cyan fluorescent protein: a sensitive pH sensor for fluorescence lifetime imaging. Anal Bioanal Chem 2013, 405(12), 3983–7.

b) Poëa-Guyon, S., Ammar, M.R., Erard, M., Amar, M., Moreau, A.W., Fossier, P., Gleize, V., Vitale, N. and Morel, N. (2013). The V-ATPase membrane domain is a sensor of pH that controls the exocytotic machinery. J Cell Biol 2013, 203(2), 283–98.

23. Porte, D.Jr. (1991). Banting lecture 1990. Beta-cells in type II diabetes mellitus. Diabetes 1991, 40, 166–180.

24. Rorsman, P. and Renström, E. (2003). Insulin granule dynamics in pancreatic beta cells. Diabetologia 2003, 46(8), 1029–45.

25. Schatz, H., Nierle, C. and Pfeiffer, E.F. (1975). (Pro-) insulin biosynthesis and release of newly synthesized (pro-) insulin from isolated islets of rat pancreas in the presence of amino acids and sulphonylureas. Eur J Clin Invest 1975, 5(6), 477–85.

26. Schmidt, W.K. and Moore, H. P. (1995). Ionic milieu controls the compartment-specific activation of pro-opiomelanocortin processing in AtT-20 cells. Mol Biol Cell 1995, 6(10), 1271–85.

27. Shibasaki, T., Takahashi, H., Miki, T., Sunaga, Y., Matsumura, K., Yamanaka, M., Zhang, C., Tamamoto, A., Satoh, T., Miyazaki, J. and Seino, S. (2007). Essential role of Epac2/Rap1 signaling in regulation of insulin granule dynamics by cAMP. Proc Natl Acad Sci USA 2007, 104, 19333–8.

28. Strasser, B., Iwaszkiewicz, J., Michielin, O. and Mayer, A. (2011). The V-ATPase proteolipid cylinder promotes the lipid-mixing stage of SNARE-dependent fusion of yeast vacuoles. EMBO J 2011, 30, 4126–41.

29. Tompkins, L.S., Nullmeyer, K.D., Murphy, S.M., Weber, C.S. and Lynch, R.M. (2002). Regulation of secretory granule pH in insulin-secreting cells. Am J Physiol Cell Physiol 2002, 283(2): C429–37.

30. Torkko, J.M., Primo, E.M., Dirkx, R., Friedrich, A., Viehrig, A., Vergari, E., Borgonovo, B., Sönmez A., Wegbrod, C., Lachnit, M., Münster C., Sica, M.P., Ermácora, M.R. and Solimena, M. (2015). Stability of proICA512/IA-2 and its targeting to insulin secretory granules require β4-sheet-mediated dimerization of its ectodomain in the endoplasmic reticulum. Mol Cell Biol 2015, 35(6), 914–27.

31. Vitavska, O., Wieczorek, H. and Merzendorfer, H. (2003). A novel role for subunit C in mediating binding of the H+-V-ATPase to the actin cytoskeleton. J Biol Chem 2003, 278(20): 18499–505.

32. Vitavska, O., Merzendorfer, H. and Wieczorek, H. (2005). The V-ATPase subunit C binds to polymeric F-actin as well as to monomeric G-actin and induces cross-linking of actin filaments. J Biol Chem 2005, 280(2): 1070–6.

33. Wilson, D.W., Lewis, M.J. and Pelham, H.R. (1993). pH-dependent binding of KDEL to its receptor in vitro. J Biol Chem 1993, 268(10): 7465–8.

34. Wu, M.M., Grabe, M., Adams, S., Tsien, R.Y., Moore, H.P. and Machen, T.E. (2001). Mechanisms of pH regulation in the regulated secretory pathway. J Biol Chem 2001, 276(35): 33027–35.

